# The neural correlates of dreaming

**DOI:** 10.1101/012443

**Authors:** F. Siclari, B. Baird, L. Perogamvros, G. Bernardi, J. LaRocque, B. Riedner, M. Boly, B. Postle, G. Tononi

## Abstract

Consciousness never fades during wake. However, if awakened from sleep, sometimes we report dreams and sometimes no experiences. Traditionally, dreaming has been identified with REM sleep, characterized by a wake-like, globally ‘activated’, high-frequency EEG. However, dreaming also occurs in NREM sleep, characterized by prominent low-frequency activity. This challenges our understanding of the neural correlates of conscious experiences in sleep. Using high-density EEG, we contrasted the presence and absence of dreaming within NREM and REM sleep. In both NREM and REM sleep, the presence of dreaming was associated with a local decrease in low-frequency activity in posterior cortical regions. High-frequency activity within these regions correlated with specific dream contents. Monitoring this posterior ‘hot zone’ predicted the presence/absence of dreaming during NREM sleep in real time, suggesting that it may constitute a core correlate of conscious experiences in sleep.

---

An ongoing stream of experiences accompanies every waking moment. Sleep is the only time in which consciousness fades under normal physiological conditions: subjects awakened from sleep, especially early in the night, report that they were not experiencing anything up to 30% of the time ^1^. At other times, subjects awakened from sleep report dreams — a stream of vivid experiences that occur despite being immobile, unresponsive, and largely disconnected from the environment. Thus, unlike wakefulness, sleep can be associated with either the presence or absence of conscious experiences. In addition, experiences in dreams can assume many forms, ranging from pure perceptual experiences to pure thought, from simple images to temporally unfolding narratives, which are often similar to awake conscious states but at times can be different in interesting ways ^2,3^.

The discovery of rapid eye movement (REM) sleep — the ‘third state of being’ besides wake and non-REM (NREM) sleep – led initially to a straightforward view of the neural correlates of dreaming ^4^: the wake-like, high-frequency, ‘activated’ EEG ^5,6^ of REM sleep was thought to be associated with the presence of dream experiences, and the low-frequency activity of NREM sleep with the absence of dreaming. However, later studies showed that up to 70% of NREM sleep awakenings yield reports of dream experiences ^1^. Conversely, in a small but consistent minority of cases, subjects deny having had any experience when awakened from REM sleep. Thus, whether one experiences something or not during sleep cannot be determined simply by assessing one’s behavioral state based on traditional EEG features or neuroimaging correlates ^7,8^.

The paradoxical occurrence of both the presence and absence of experiences within the same behavioral state of sleep, and across two very different kinds of sleep (NREM and REM) challenges our current understanding of the neural correlates of conscious experience in sleep.

Here we investigate the neural correlates of dreaming using a within state paradigm, in both NREM and REM sleep, by performing serial awakenings of subjects recorded throughout the night with high-density EEG (256 channels)^9^. The results highlight a posterior cortical ‘hot zone’ where a local decrease in low-frequency EEG activity during both NREM and REM sleep is associated with reports of experiences upon awakening, whereas a local increase in low-frequency activity is associated with the absence of experience. These results hold for both a large group of naïve subjects and a smaller group of individuals trained in dream reporting. We then show, in a separate group of subjects, that it is possible to predict whether an individual will report having dreamt or not in the course of a NREM episode by real-time EEG monitoring of this posterior hot zone. Finally, we show that the location of high-frequency EEG activity during dreams correlates with specific dream contents, such as thoughts, perceptions, faces, spatial setting, movement, and speech.

## Results

Using a serial awakening paradigm ^10^, participants were awakened throughout the night and were asked to report whether, just before the awakening, they had been experiencing anything (dreaming experience, DE), experiencing something but could not remember the content (dreaming experience without recall of content, DEWR), or not experiencing anything (no experience, NE, see *Online Methods*). If subjects reported a DE they were asked to describe its most recent content (‘the last thing going through your mind prior to the alarm sound’), and to rate it on a scale ranging from exclusively thought-like (thinking or reasoning, with no sensory content) to exclusively perceptual (vivid sensory content, without thinking or reasoning)^11^. Finally, they had to estimate the duration of the most recent DE, and to report whether it contained specific categories of content, including faces, a spatial setting, movement, and speech (examples of reports are presented in Supplementary table S1).

We studied two different populations in separate experiments. In experiment 1 we studied a large group of subjects who underwent few awakenings (N=32, 233 awakenings), while in experiment 2, we investigated a smaller group of trained subjects who underwent many awakenings (N=7, 815 awakenings) (see Supplementary table S2). Finally, in a third experiment, we tested whether, based on the results of the first two experiments, we could predict the presence or absence of dreaming in real time at the single trial level (N=7, 84 awakenings).

### Dreaming experience vs no experience: low-frequency power

First, we sought to determine whether and how low-frequency activity (1-4 Hz) changes between the presence and absence of conscious experience during sleep. EEG slow frequencies between 1 and 4 Hz during sleep are associated with neuronal down-states and bistability ^12^, which prevent the emergence of stable causal interactions among cortical areas ^13^, and have been linked to the loss of consciousness ^14,15^. For this analysis we focused on the first group of subjects (experiment 1) because of the large number of subjects, and on N2 sleep, since the number of DE and NE reports in this stage were well balanced. By computing power spectral density at the source level within the 1-4 Hz frequency band we found that reports of DE, compared to reports of NE, were preceded by decreased low-frequency power. This decrease was restricted to a bilateral parieto-occipital region, encompassing the medial and lateral occipital lobe and extending superiorly to the precuneus and posterior cingulate gyrus (Fig. 1A). Experiment 2, in which awakenings were carried out both in N2 and N3 stage sleep, confirmed these results (Supplementary Figure 1).

**Figure 1.**
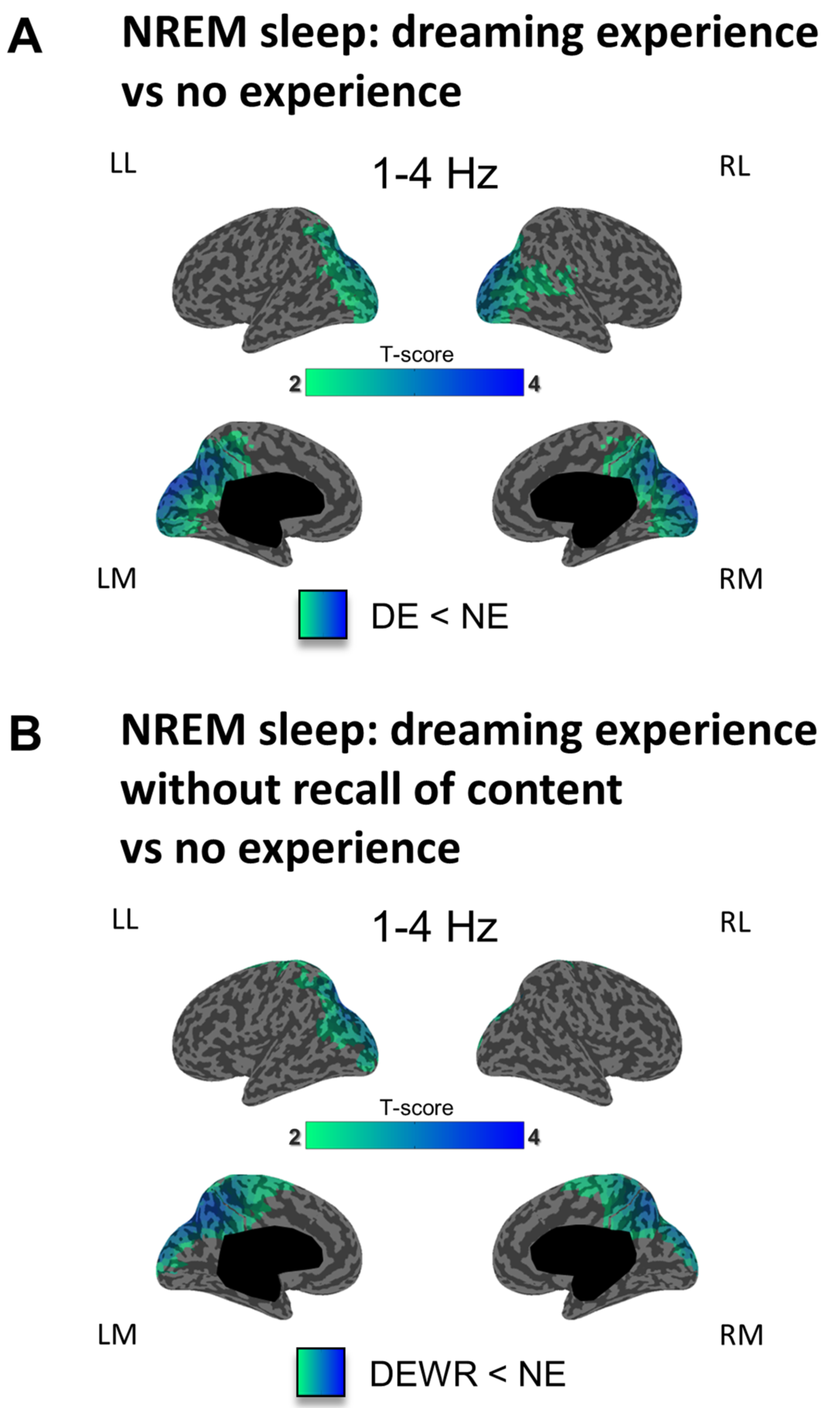
Dreaming experience vs. no experience in NREM sleep (low-frequency power). A. Inflated cortical maps illustrating the topographical distribution of t-values for the contrast between DEs and NEs at the source level for low-frequency power (1–4 Hz) in NREM sleep (last 20 seconds before the awakening). Only significant differences at the p<0.05 level, obtained after correction for multiple comparisons are shown (two-tailed, paired t-tests, 32 subjects, t(31) > 2.04). B. Same as A for the contrast between NE and DEWR (two-tailed, paired t-tests, 20 subjects, t(19) > 2.09).

We then compared instances in which the subjects reported that they had dreaming experiences, but were not able to recall the content of the experience (DEWR), with instances in which they reported being unconscious (i.e., having no experiences, NE). Reports of DEWR, contrasted to reports of NE, were again preceded by decreased low-frequency power in a parieto-occipital region highlighted by the DE/NE contrast (Fig. 1B, see Supplementary Figure 2 for a conjunction map of the two contrasts). A contrast between DE and DEWR in the low-frequency range did not yield significant differences. Thus, decreased low-frequency power in this posterior cortical zone was correlated with the occurrence of dreaming experiences, irrespective of the ability to remember the contents of the experience.

Next, we asked whether the contrast between DE and NE would produce similar results also during REM sleep, a state that is substantially different from NREM sleep in terms of EEG signatures (slow waves and spindles in NREM sleep, low-voltage fast activity in REM sleep ^4^), neural activity (widespread bistability between down and up-states in NREM sleep, mostly tonic depolarization in REM sleep ^6^), neuromodulation (high acetylcholine in REM sleep, low in NREM sleep ^16^), and regional activations^7^. Because of the rarity of NE reports in REM sleep (Supplementary table S2), for this analysis we combined the data of experiments 1 and 2. Despite the notable physiological differences between NREM and REM sleep states, we again found that brain activity associated with DE, as compared to NE, had reduced power in the 1-4 Hz band in a parieto-occipital region (Fig. 2A), which largely overlapped with the topography observed in NREM sleep (Fig. 2B). Thus, a decrease in low-frequency power in a posterior cortical ‘hot zone’ is highlighted when dreaming experiences are contrasted with the absence of experience during sleep, irrespective of the ability to recall the experience and of behavioral state (NREM vs. REM sleep).

**Figure 2.**
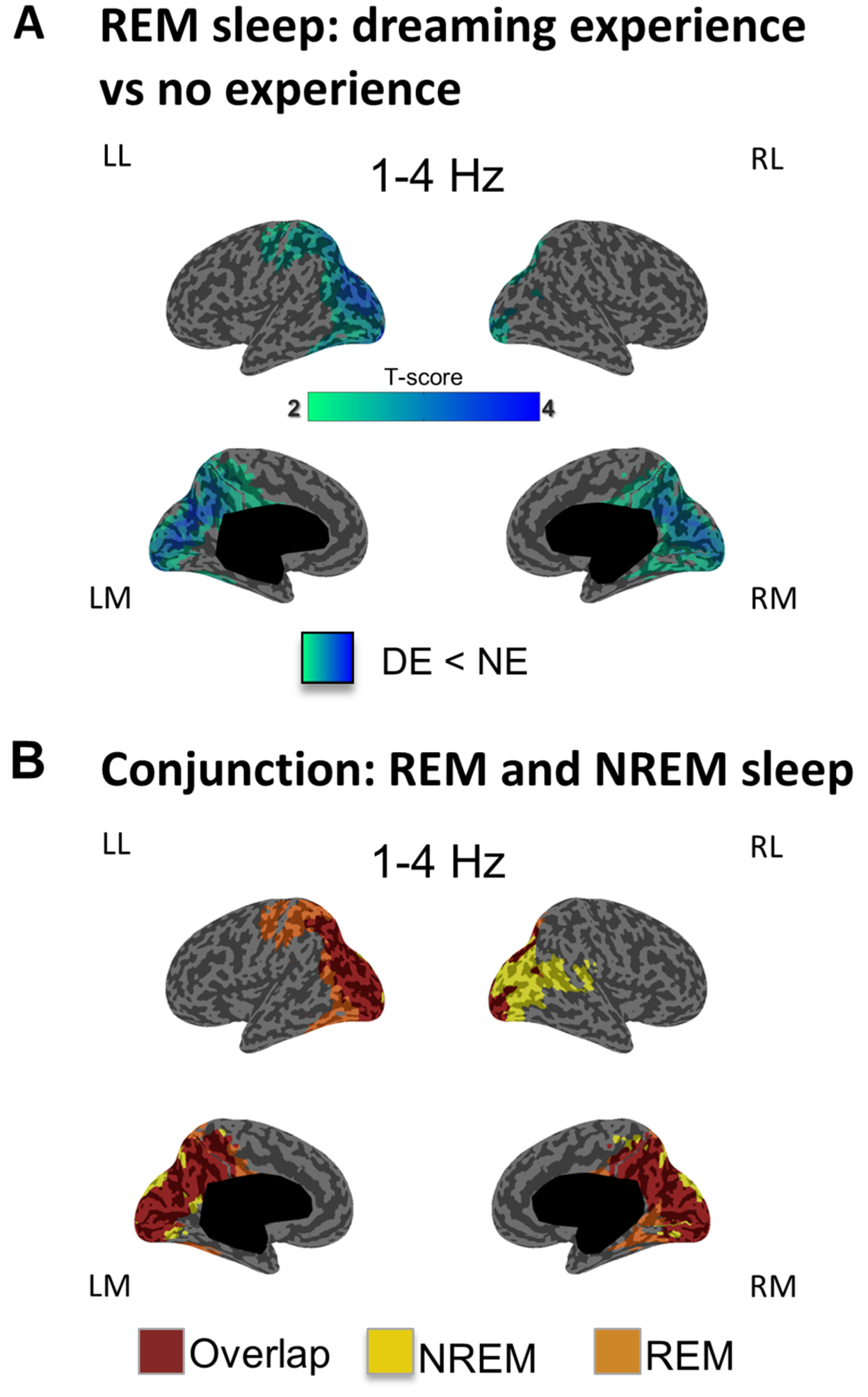
Dreaming experience vs. no experience in REM sleep (low-frequency power). **A.** Inflated cortical maps illustrating the topographical distribution of t-values for the contrast between DEs and NEs at the source level for low-frequency (1–4 Hz) power in REM sleep (20 seconds before the awakening). Only significant differences at the p<0.05 level, obtained after correction for multiple comparisons are shown (two-tailed, paired t-tests, 10 subjects, t(9)>2.26). **B**. Conjunction maps showing the differences and overlap compared to the same contrast performed in NREM sleep.

### Dreaming experience vs no experience: high-frequency power

We then investigated differences in high-frequency band power (20-50 Hz), since high EEG frequencies often reflect high rates of neuronal firing ^17,18^ and may thus indicate brain regions that show increased activity. In NREM sleep, DE, compared to NE, was associated with increased high-frequency power in the same parieto-occipital region that showed reduced low frequency power. However, differences in high-frequency power extended superiorly and anteriorly to parts of the lateral frontal cortex and the temporal lobes (Fig. 3A). DE with recall of content, compared to DEWR, were associated with higher high-frequency power in medial and lateral frontoparietal areas (Fig. 3B). No differences in high frequency power were found when comparing DEWR with NE. Finally, in REM sleep DEs compared to NEs were associated with increased high-frequency (25-50 Hz) power in frontal and temporal regions (Fig. 3C).

**Figure 3.**
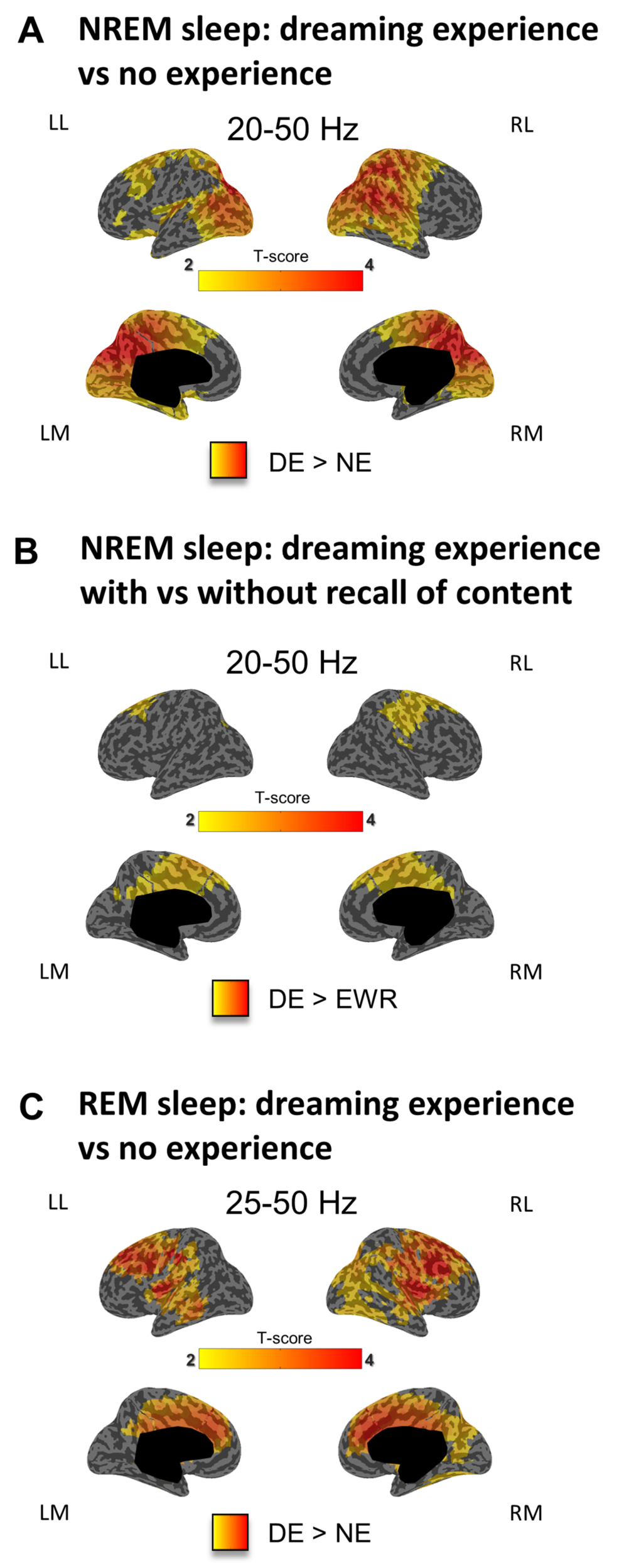
Dreaming experience vs. no experience in NREM and REM sleep (high-frequency power). **A.** Inflated cortical maps illustrating the topographical distribution of t-values for the contrast between DEs and NEs at the source level for high-frequency power (20–50 Hz) in NREM sleep (last 20 seconds before the awakening). Only significant differences at the p<0.05 level, obtained after correction for multiple comparisons are shown (two-tailed, paired t-tests, 32 subjects, t(31) > 2.04). ***B***. Same as A for the contrast between DE and DEWR in NREM sleep (two-tailed, paired t-tests, 20 subjects, t(19) > 2.09) ***C***. same as A for high-frequency power (25-50 Hz) in REM sleep (two-tailed, paired t-tests, 10 subjects, t(9) > 2.26).

### Other analyses

To test whether our results were characterized by a significant hemispheric lateralization, we computed a repeated-measure-ANOVA with hemisphere and type of report (DE, DEWR, NDE) as within-subject factors for all the contrasts (See ‘Online Methods’ and Supplementary table 5). We did not find significant interactions between the type of report and hemisphere, except for a barely significant right-sided lateralization (p= 0.046) for the high-frequency power contrast between DE and NE in REM sleep.

Absolute power values for DE and NE in the posterior hot zone are shown in Supplementary Figure 3. It should be noted that absolute low-frequency power values are higher in NREM sleep compared to REM sleep, although differences are comparable and significant in both sleep stages. It is likely that the absolute values in NREM sleep, although obtained from a region of interest after source modeling the signal, are nevertheless influenced by high-amplitude slow waves in other parts of the cortex in NREM sleep. Indeed, source modeling strongly reduces but does not completely eliminate the effects of volume conduction. Moreover, regularization methods commonly applied to reduce the influence of noise in EEG data introduce a ‘smoothing’ effect of the EEG sources. Finally, the effect of slow wave propagation, which occurs on a scale of tens/hundreds of milliseconds is not affected by source modeling approaches and may contribute to the relatively high level of low-frequency activity all over the cortical mantle and in brain areas that are not generating slow waves.

Given the age variability in our sample, an additional ROI-based repeated measures ANCOVA was performed in the significant clusters using this parameter as nuisance covariate. A significant main effect of report type (DE, NE) was confirmed for both low-frequency (*p*=0.003) and high-frequency activity (*p*=0.005), while no significant effects of age emerged (*p* > 0.05).

### Dream content in REM sleep

We further investigated whether differences in high-frequency power could distinguish specific contents of DEs. For these analyses, we focused on REM sleep awakenings in trained subjects (experiment 2), because there were many intra-subject trials, reports were more detailed ^10^ and subjects were more confident about their answers concerning specific contents.

Since dreams can differ greatly along the dimension ‘perception vs. thought’^11^, subjects had been asked to rate their DEs accordingly. We found that high-frequency activity correlated with the thinking dimension in frontal brain regions and with the perceiving dimension in parietal, occipital and temporal areas (r>0.14, Fig. 4A). This anterior-posterior gradient suggests that in dreams, just like in wakefulness, anterior cortical regions mediate thought-like experiences while posterior regions are involved in perceptual aspects of the experience.

**Figure 4.**
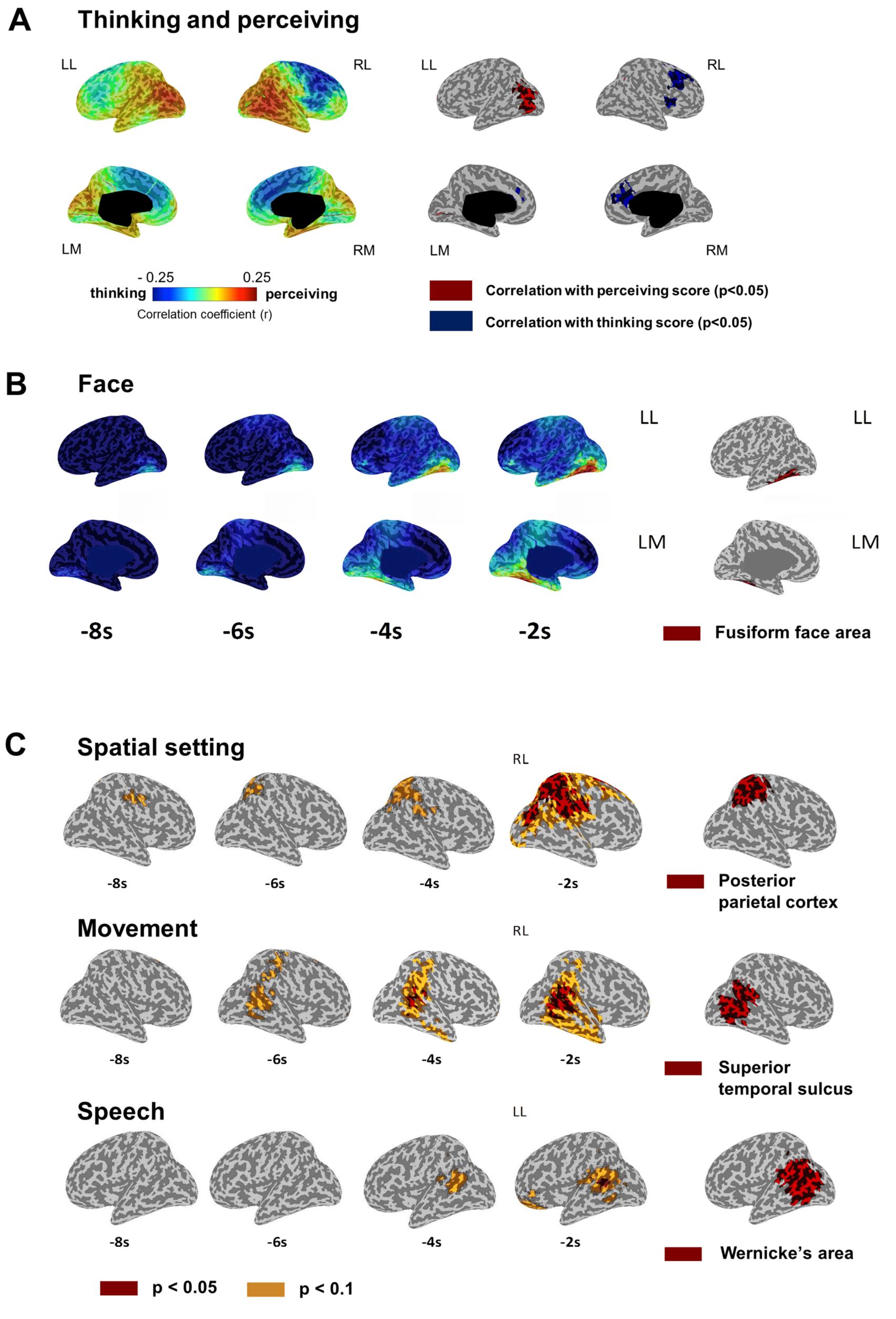
The content of dream experiences in REM sleep. **A.** Correlation between the thinking/perceiving score and 25–50 Hz power (last 8s, 7 subjects). *Left*: mean Spearman rank correlation coefficients (7 subjects). *Right:* significant voxels p<0.05 (one-tailed permutation test, r>0.14). **B.** *Left:* 25–50 Hz power differences (DE with face *minus* DE without face). ROI contrast R FFA p=0.023; one-tailed paired t-test (7 subjects). *Right*: fusiform face area^19^ (red). **C. Upper row:** 25–50 Hz average power differences between DEs with and without a spatial setting (6 subjects, t(5) > 2.57), *Right*: right posterior parietal cortex^21^, Middle row: movement vs. no movement (7 subjects, t(6) > 2.45). Right: superior temporal sulcus ^22^, Bottom row: speech vs. no speech (7 subjects, t(6) > 2.45). Right: Wernicke’s area ^24^. Two-tailed paired t-tests. p<0.05 (red) and p<0.01 (yellow).

We also examined specific perceptual categories that were experienced during sleep. As shown in Fig. 4B, DEs containing faces, compared to DEs without faces, were associated with increased high-frequency activity in a circumscribed temporo-occipital region that closely matched the fusiform face area (FFA). A region of interest (ROI) analysis^19^ confirmed that high-frequency activity in the FFA on the right side was significantly higher for DEs containing faces (p=0.023, paired one-tailed t-test) compared to DEs without faces, consistent with the specialization of this area in the perception of faces during wakefulness ^19^. Subjects were also asked to determine whether the setting of the most recent E was indoors, outdoors, or could not be specified. We found that DEs with a definite spatial setting were associated with increased high-frequency activity in the right posterior parietal cortex (rPPC) (Fig. 4C), an area involved in spatial perception and visuo-spatial attention ^20^, whose lesion can cause spatial neglect. This finding was confirmed by a ROI analysis ^21^ (p=0.023, paired one-tailed t-test). Furthermore, DEs in which subjects reported the sense of moving in the dream were associated with increased high-frequency activity in a region surrounding the right superior temporal sulcus (Fig. 4C), an area involved in the perception of biological motion ^22^ and viewing body movements ^23^. A ROI analysis ^22^ confirmed this finding (p=0.029, paired one-tailed t-test). Finally, as shown in Fig. 4C, DEs containing speech were associated with increased high frequency activity over a left posterior temporal region corresponding to Wernicke’s area, as confirmed by a ROI analysis^24^ (p=0.048, paired one-tailed t-test).

### Real-time prediction of dreaming in NREM sleep

If a decrease in low-frequency EEG power and an increase in high-frequency power in a posterior hot zone represents a reliable neural correlate of dreaming, it should be possible to predict whether a person is dreaming or not in real time by detecting ongoing local EEG activations. Moreover, this should be the case even though the global EEG, as well as the local EEG in anterior areas, is characterized by slow-wave activity typical of NREM sleep. To test this prediction, seven additional subjects slept for three consecutive nights in the sleep laboratory while high-density EEG was continuously recorded. Based on the findings of Experiments 1 and 2, on the experimental (second and third) nights participants were awoken from NREM sleep when their neural activity surpassed a bispectral threshold in low-frequency (0.5-4.5 Hz) and high-frequency (18-25 Hz) band power from an electrode cluster over the posterior hot zone (Fig. 5A; see *Online Methods*; the 18-25 Hz range was used because it captures EEG activations while avoiding artifacts from eye movements and other sources during online EEG monitoring, see also *Online Methods*). In total, participants were awakened 84 times (mean: 12.0 ± 3.4; Supplementary table S3). After removal of trials containing artifact (see *Online Methods*), there remained 36 awakenings initiated by the algorithm with a DE prediction, out of which 33 were accurate (DE mean accuracy = 91.6%, *p*=0.00001), and 26 awakenings with a NE prediction, out of which 21 were accurate (NE mean accuracy = 80.7%, *p*=0.0003); total prediction accuracy across all states was 87% (Fig. 5B). No interaction was observed between DE prediction and time of night (*F*_(1,6)_=1.35, *p*=0.29). Confirming significant differences in EEG power in DE compared to NE trials in the prediction ROI, we observed significantly lower low-frequency activity (*p*=0.001) in DE (10.17±2.5 [mean ± SD]) compared to NE (103.47±38.16) and significantly higher high-frequency activity (*p*=0.001) in DE (0.14±0.03) compared to NE (0.07±0.02) prediction trials (Fig. 5C). As shown in Figure 5D and Supplementary table S4, DE had a higher high/low frequency band power ratio compared to NE in cortical regions including bilateral occipital cortex, precuneus, right superior parietal lobule (SPL), right precentral gyrus, left superior and middle temporal sulcus (STS), right lingual gyrus and left inferior frontal gyrus (IFG) (*p* < 0.001, FDR cluster corrected).

**Figure 5.**
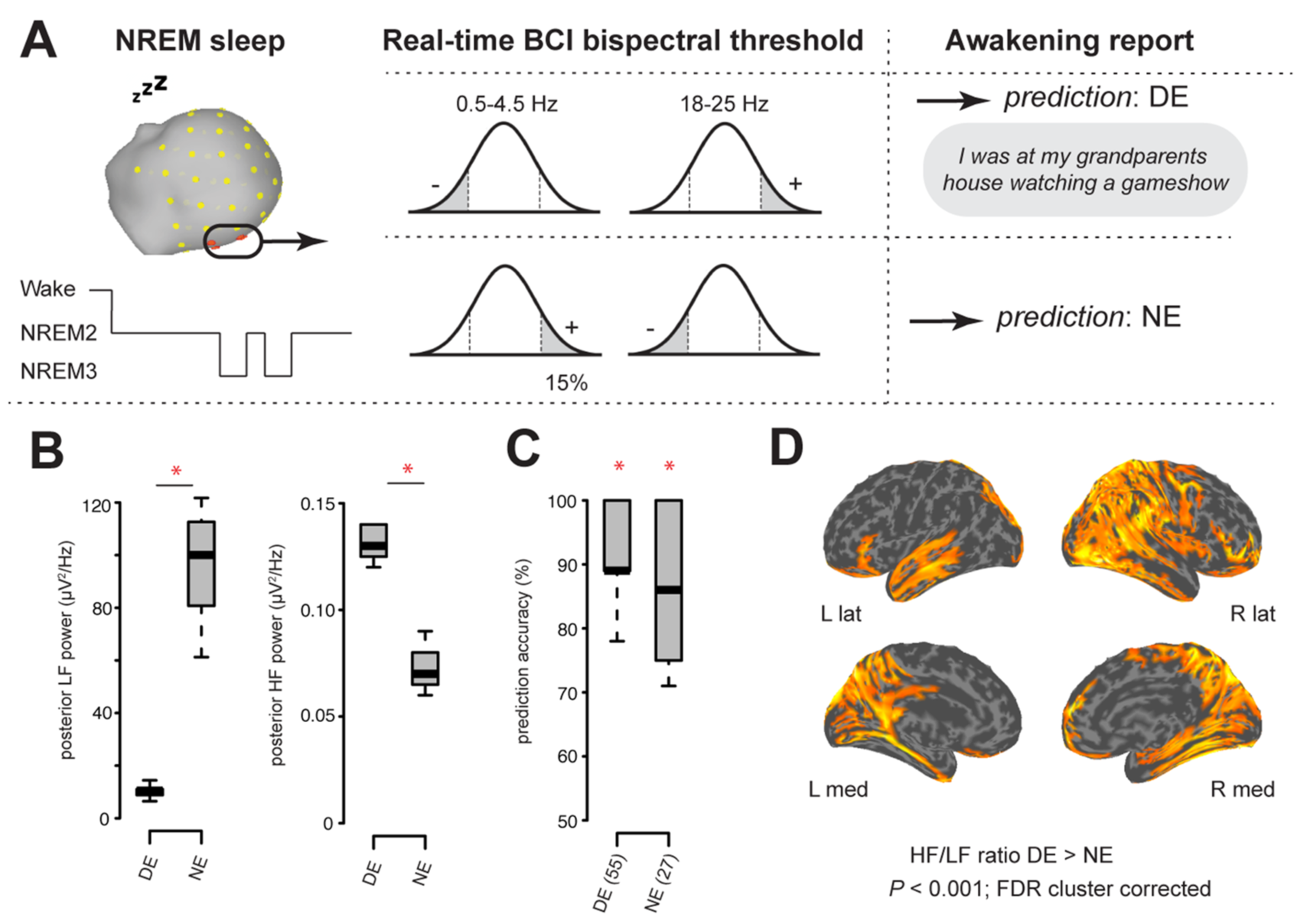
Real-time prediction of dream experience. **A.** Seven participants were awoken from NREM when neural activity surpassed a bispectral threshold in low-frequency (LF; 0.5–4.5 Hz) and high-frequency (HF; 18–25 Hz) power over the posterior hot zone. **B.** DE trials had significantly lower LF activity and significantly higher HF activity in DE compared to NE prediction trials. **C.** Prediction accuracy for E (55) and NE (27) trials. **D.** DE had a higher HF/LF power ratio compared to NE in cortical regions including bilateral occipital, medial and lateral parietal, medial temporal and inferior frontal cortex (p<0.001, FDR cluster corrected).

## Discussion

The three experiments reported here highlight a parieto-occipital ‘hot zone’ as the neural correlate of dreaming. When this posterior hot zone showed a decrease in low-frequency EEG activity – traditionally known as ‘EEG activation’^5,6^ – subjects reported upon awakening that they did have dream experiences. By contrast, when low-frequency EEG activity increased in the same area, subjects reported that they had been unconscious. This result held true whether the subjects had been in NREM sleep or in REM sleep, irrespective of the dominant EEG activity over other cortical regions.

The influential notion that dreaming is virtually synonymous with REM sleep has dominated neuroimaging work for the past several decades ^7^, despite the increasing number of studies suggesting that dream reports could be obtained from every stage of NREM sleep, albeit less often ^25^. The frequent conflation of REM sleep with dreaming was eased by the marked similarities between the overall EEG of REM sleep and that of waking consciousness, and little attention was given to the rare reports of REM sleep awakenings that yielded no reports. Conversely, reports of dreaming from NREM sleep awakenings were hard to reconcile with EEG patterns characterized by slow wave activity – the opposite of EEG activation ^5^. Indeed, it was suggested that reports of dreaming after NREM awakenings were due to ‘covert’ REM sleep ^26^ ^26^, or even that neural activity in the cortex was unrelated to the presence or absence of consciousness ^27^. The present results resolve these paradoxes by showing that in both REM and NREM sleep, dreaming, and more generally, conscious experience during sleep, requires a localized activation of a posterior hot zone, irrespective of the EEG in the rest of the cortex. This explains why dreaming can occur in two behavioral states that differ radically in terms of global EEG signatures.

Of course, since the posterior hot zone was highlighted by averaging EEG recordings preceding many awakenings from many subjects, the full neural correlates of experiences ins sleep could occasionally include additional brain regions. For example, dreams that include insight into the state (so-called lucid dreams) or dreams in which an individual has some degree of control over the content of the dream may recruit additional areas of frontal-parietal cortex ^28^ ^29^. Conversely, we cannot rule out that the posterior hot zone may include, besides brain regions supporting experiences, areas that might support unconscious aspects of dream generation. However, many studies contrasting perceiving or not specific sensory stimuli during wakefulness (such as seeing/not seeing a face, an object, a place, a movement, and so on) also point to regions within the posterior hot zone ^30^. EEG source modeling indicates that, in addition to low and high-level sensory areas (primarily visual), the posterior hot zone prominently includes precuneus, posterior cingulate, and retrosplenial cortices: areas on the medial surface of the brain whose electrical stimulation can induce a feeling of being ‘in a parallel world’^31^, disconnected from the environment in a ‘dream-like state’^32^, and whose involvement in multisensory integration is well-suited to support the virtual simulation of a world ^33^ and ‘immersive spatio-temporal hallucinations’^9^ that characterize dreams.

The posterior hot zone was also highlighted by the contrast between the absence of experiences and experiences without recall of specific contents of the experience. This finding further suggests that the activation of this posterior hot zone is a marker of experiences themselves, rather than of the recall of experiences. Indeed, both lesion studies and neuroimaging results indicate that specific contents of experiences are supported by posterior cortical areas ^30,34^,^35^, while a different set of brain areas are involved in encoding and storing memories and in directing attention ^36,37^. The results we obtained were also virtually identical in naïve subjects (Experiment 1) and in trained subjects (Experiment 2), suggesting that they characterize a core correlate of dreaming, rather than some other cognitive function.

In our experiments conscious experiences were consistently associated with low activity in the delta range (1–4 Hz) and the absence of experience with high delta activity in the posterior hot zone. During NREM sleep, EEG slow waves in the delta range reflect alternations between periods of depolarization (up-states), during which neurons can fire, and periods of hyperpolarization (down-states) during which neurons cease firing ^12^. This bistability of membrane potential causes the breakdown of stable causal interactions among cortical areas ^13^, which are thought to be necessary for sustaining conscious experiences ^14,15^. Recently, the occurrence of local slow waves has also been discovered during REM sleep, especially in the supragranular layers of primary cortical areas ^38^, adding to the evidence that neuronal activity during sleep can have a local component ^12^. A global slowing of EEG activity into the delta range had been associated with loss of consciousness in many previous studies ^14,15,39,40^ However, there are several situations in which subjects can be conscious and have experiences despite the presence of local and global slow wave activity ^30,41,42^, including of course NREM sleep itself, where global slow wave activity is present despite the frequent occurrence of dreaming. The present findings suggest that future work in patients with disorders of consciousness, epileptic seizures, or under general anesthesia should examine the level of EEG activation in the posterior hot zone to uncover the possible occurrence of consciousness despite the loss of responsiveness.

In addition to differences in EEG low-frequency activity in the posterior hot zone, we found that this region also showed increased high-frequency activity when subjects reported dreaming experiences during sleep. High-frequency activity usually reflects increased neuronal firing ^17^. A plausible interpretation of this finding is that, as long as the posterior hot zone is not in a low-frequency mode that prevents experiences, the specific contents of experiences when dreaming are dictated by which specific groups of neurons fire strongly and which do not, just as in wake. Consistent with this, we found that specific contents of a subject’s REM sleep dream – such as thoughts, perceptions, faces, places, movement, and speech – are associated with increased high-frequency EEG activity in specific cortical areas, which correspond closely to those engaged during waking perception of the same contents. In a recent fMRI study, a pattern classifier trained while subjects perceived natural images during wakefulness ^43^ was able to successfully decode specific content categories reported by subjects awakened at the transition between wakefulness and sleep. Our results with high-density EEG extend these findings beyond falling asleep to periods of consolidated sleep, providing further evidence that dream reports reflect conscious experience during sleep rather than confabulations upon awakening ^44,45^.

Increased high-frequency activity associated with reports of dreaming experiences were also observed in areas that went beyond the posterior hot zone. Thus, increased high-frequency activity extended to some portions of prefrontal cortex during REM sleep dreaming, especially for thought-like experiences. Moreover, a frontal cluster (Fig 3B) showed increase high-frequency activity during NREM sleep when contrasting experiences with and without recall of content, suggesting that this area may be important for recall of the content of the experience. Neuronal activity in the gamma range was initially linked to consciousness, but later studies showed that consciousness and gamma activity can be dissociated ^30^. For example, one can be aware of a stimulus whether or not it triggers gamma activity ^46^, and gamma activity can be strong during general anesthesia in the presence of low frequency oscillations ^47^. However, recent studies have reported that gamma-band transcranial alternating current stimulation is associated with higher scores on the revised coma scale recovery scale (CRS-R) ^48^ or self-awareness during REM sleep dreams ^49^, suggesting that high-frequency activity may indeed influence functions related to cognition or awareness. It is thus an open question whether high-frequency activity in a subset of prefrontal regions may contribute some specific contents to experience ^30^, such as conscious thought or mind-wandering states ^50^, emotional/motivational contents ^51^, metacognition ^52^, or insight into one’s state (i.e., lucid dreaming)^28,29^. Alternatively, high-frequency activity in prefrontal regions may mediate cognitive functions that are unconscious but affect experience indirectly, such as the encoding and storing of long-term memories, the deployment of attention, and the planning and execution of tasks.

Finally, this study suggests that dreaming may constitute a valuable model for the study of consciousness with implications beyond sleep. First, the direct comparison between consciousness (the presence of experience) and unconsciousness (the absence of experience) within the same behavioral state avoids confounds that limit the interpretation of between-states paradigms. These confounds arise because, in studies contrasting wakefulness with sleep, anesthesia, or coma, many other aspects of brain function besides consciousness change along with behavioral state changes^53^. Second, during sleep subjects are largely disconnected from the environment on both the input and output side, meaning that in the majority of cases, sensory stimuli generally do not influence the content of experience^2^. Third, during sleep subjects are not performing an experimental task, avoiding confounds caused by stimulus processing, response preparation, and execution ^53^.

An influential notion derived from neuroimaging studies contrasting wake with NREM sleep, anesthesia, or coma is that the neural correlates of consciousness would correspond to a broad fronto-parietal network ^54^. This notion has been reinforced by findings indicating that the same fronto-parietal network is activated when contrasting brain responses to the same stimuli when subjects reported seeing them or not ^55^. However, lesion studies suggest that consciousness persists after widespread frontal lesions ^30,56^ and recent work controlling for confounding factors related to stimulus processing, attention, and task performance have cast doubt on a central role of this fronto-parietal network in supporting consciousness ^30,57^,^58^. By exploiting a within state, no stimulus, no task paradigm, our findings indicate that the neural correlate of consciousness, rather than involving the broad lateral fronto-parietal network, is restricted to a posterior hot zone.

## Online Methods

### Experiments 1 and 2

#### Procedure

The procedure used in this study has been described in detail in a previous publication ^10^. Awakenings in the sleep laboratory were performed at pseudorandom intervals, irrespective of sleep stage, using a computerized sound that lasted 1.5 sec, which was administered through E-Prime (Psychology Software Tools, Pittsburgh, PA). Subjects were instructed to signal that they had heard the alarm sound and to lie quietly on their back. They then underwent a structured interview via intercom about their mental activity that lasted between 20 sec and 3.5 min, depending on whether the subject reported a conscious experience and had to answer additional questions related to the content. Signed informed consent was obtained from all participants before the experiment, and ethical approval for the study was obtained from the University of Wisconsin-Madison Institutional Review Board.

#### Study participants experiment 1

Thirty-two healthy subjects were included in this experiment (12 males, age 46 ± 13.3 years, 24–65 [mean ± SD, range]) that were selected from a group of 69 subjects participating in a larger research project in our laboratory. Subjects were randomly selected from the subset of all participants that had at least one DE and NE report in NREM sleep during the same night. Among the 240 awakenings performed in these 32 subjects, 5 had to be excluded for technical problems and 2 because subjects were too somnolent upon awakening to answer questions reliably. Comparisons between DEWR-NE conditions and DE-DEWR conditions were performed in the subset of these participants (N=20) that had all three types (DE, DEWR, NE) of report during the same night. In REM sleep, only 6 out of the 69 participants presented both DE and NE. They were all included in the analysis.

#### Study participants experiment 2

Seven healthy subjects were included in this experiment (3 males, age 31 ± 8.8 years, 21–47 [mean ± SD, range]). None of the subjects in this experiment participated in experiments 1 or 3. The behavioral results of 6 of these 7 subjects (i.e., dream recall rates and characteristics of dreams across sleep stages), but not the EEG results, have been described in a previous publication ^10^. Data from experiments 1 and 3 has not been reported previously.

Study participants received comprehensive explanations regarding the questionnaire used in the experiment, and filled it in at home every morning upon awakening for two weeks before the first study night. During this training phase experimenters communicated repeatedly with the subjects to ensure that they were proceeding with the training and to answer any questions the subjects had regarding the questionnaire. Between five and ten overnight recordings were performed for each participant, with a maximum of three consecutive nights. To increase the number of awakenings in REM sleep with respect to NREM sleep, two additional nights were scheduled for three of the subjects, in which awakenings were only performed in REM sleep. The total number of nights was distributed as follows among subjects: S1:10 nights, S2:8 nights, S3:6 nights, S4:5 nights, S5:8 nights, S6:6 nights, S7:8 nights. Overall, 836 awakenings were performed, of which 5 had to be excluded because of technical problems, and 16 because subjects were too somnolent upon awakening to answer questions reliably.

#### Sleep recordings

Recordings were made at the University of Wisconsin Center for Sleep Medicine and Sleep Research (WisconsinSleep). They were performed using a 256-channel high-density EEG (hd-EEG) system (Electrical Geodesics, Inc., Eugene, Ore.). Four of the 256 electrodes placed at the outer canthi of the eyes were used to monitor eye movements; additional polysomnography channels were used to record submental electromyography. Sleep scoring was performed over 30s epochs according to standard criteria ^59^. Recordings lasted approximately 8h and were performed with EEG nets that are validated for long-term monitoring for up to 24 hours. Impedances were checked at the beginning and end of each recording. An experimenter continuously evaluated the signal during the night, and repositioned electrodes or filled them with additional gel upon noticing artifacts in the signal. As described below, a thorough artifact-removal procedure was applied offline.

#### Preprocessing of data experiment 1

The signal corresponding to the 2 mins preceding each awakening was extracted and considered for analysis. The EEG signal was sampled at 500 Hz and off-line band-pass filtered between 1 and 50 Hz for the NREM data and between 0.3 and 50 Hz for the REM data, in order to be merged with the REM data of Experiment 2. The NREM EEG data was high-pass filtered at 1Hz instead of lower frequencies, as there were strong sweating artifacts in some of the participants that caused intermittent high-amplitude slow frequency oscillatory activity around 0.3 Hz. REM data did not contain any sweating artifacts and was high-pass filtered at 0.3Hz. Channels containing artifactual activity were visually identified and replaced with data interpolated from nearby channels using spherical splines (NetStation, Electrical Geodesic Inc.). To remove ocular, muscular, and electrocardiograph artifacts we performed Independent Component Analysis (ICA) using EEGLAB routines ^60^. Only ICA components with specific activity patterns and component maps characteristic of artifactual activity were removed ^61^.

#### Preprocessing of data experiment 2

The signal corresponding to the 2 mins preceding each awakening was extracted and considered for analysis. The EEG signal was sampled at 500 Hz and off-line band-pass filtered between 0.3 and 50 Hz for both NREM and REM data. Channels containing artifactual activity were visually identified and replaced with data interpolated from nearby channels using spherical splines (NetStation, Electrical Geodesic Inc.). To remove ocular, muscular, and electrocardiograph artifacts we performed Independent Component Analysis (ICA) using EEGLAB routines ^60^. Only ICA components with specific activity patterns and component maps characteristic of artifactual activity were removed ^61^.

#### Signal analysis

##### Source localization

The previously cleaned, filtered and average-referenced EEG signal corresponding to the 20s before the awakening was extracted and analyzed at the source level. Source modelling was performed using the GeoSource software (Electrical Geodesics, Inc., Eugene, Ore.). A 4-shell head model based on the Montreal Neurological Institute (MNI) atlas and a standard coregistered set of electrode positions were used to construct the forward model. The source space was restricted to 2447 dipoles that were distributed over 7x7x7 mm cortical voxels. The inverse matrix was computed using the standardized low-resolution brain electromagnetic tomography (sLORETA) constraint ^62^. A Tikhonov regularization procedure (λ=10^−1^) was applied to account for the variability in the signal-to-noise ratio ^62^. We computed spectral power density using the Welch’s modified periodogram method (implemented with the *pwelch* function in MATLAB (The Math Works Inc, Natick, MA) in 2s Hamming windows (8 segments, 50% overlap) to decompose the source signals into frequency bands of interest.

##### Statistical Analysis

Statistical analyses were carried out in MATLAB. To compare brain activity between DE and NE, the source-modelled signal was averaged within the time periods and frequency bands of interest. We then averaged the signal associated with DEs and NEs within each subject and for each frequency band and stage (REM and NREM sleep) separately. Whole-brain group level analyses on average absolute power values were performed separately for each frequency range using a within-subjects design, and corrected for multiple comparisons using Statistical nonparametric mapping (SnPM)^63^.

For analyses of content (faces, spatial setting, movement, speech), we used a more liberal threshold (p<0.05, uncorrected, two-tailed paired t-tests), given that there was a strong a priori hypothesis about the direction of the contrast and activations were expected in restricted cortical regions. We first verified that the data were normally distributed using the Lilliefors modification of the Kolmogorov-Smirnov test. For all planned comparisons (face, setting, movement, speech), a normal distribution was confirmed for the large majority of voxels (range 81.2-95.0%). Given the relative robustness of parametric t-tests to violations of normality, and the relatively small percentage of voxels breaking this assumption, potential differences were assessed using two-tailed paired t-tests (p<0.05).

Additional region of interest (ROI) based analyses were performed as follows: ROIs were defined as 15mm radius spheres centered on coordinates (ex. fusiform face area) obtained from previous imaging studies. Then, high-frequency power values (25–50 Hz) were averaged within these ROIs and compared between conditions (i.e., face vs no face) using one-sided paired t-tests. Larger ROIs (25mm spheres) was used for the figures (Figure 3) for better visibility.

To investigate the correlation between the thinking/perceiving score and high frequency power (25–50 Hz), we calculated the Spearman rank correlation coefficient for each subject and voxel, and then averaged the resulting coefficients across subjects to obtain group-level correlation maps (Fig. S3). Statistical significance was determined using a permutation-based approach. Specifically, null distributions were generated for each voxel by randomly shuffling the two variables of interest (thinking/perceiving score and high-frequency power) in each subject and recalculating the group-level mean correlation (n=1000). A voxel-wise parametric one-sided t test was subsequently performed (p < 0.05).

To assess lateralization of findings, for each contrast we evaluated the potential difference in the number of voxels showing a significant effect between the two hemispheres by calculating a lateralization index (LI) as described in ^64^. To test for a statistically significant difference between the level of activity observed in symmetrical areas across the two hemispheres, we calculated the relative difference in power spectral density (PSD) between two symmetrical ROIs including the voxels that showed a significant effect in the hemisphere characterized the stronger LI, but not in the contralateral side. In particular, we explored the potential interaction between hemisphere (Left, Right) and type of report (DE, NDE, DEWR) using a repeated measure ANOVA.

### Experiment 3

#### Study Participants Experiment 3

Seven additional participants (4 men, 3 women; age range = 19–31 years) not included in Experiments 1 or 2 were recruited for Experiment 3. All participants were right-handed and had no history of neurological disorder. Signed informed consent was obtained from all participants before the experiment, and ethical approval for the study was obtained from the University of Wisconsin-Madison Institutional Review Board.

#### Sleep Recordings

Hd-EEG was continuously recorded overnight using a 64-electrode EEG cap (EasyCap, Herrsching, Germany) with equidistantly spaced sintered Ag/AgCl ring electrodes. The signal was amplified using BrainAmp MR Plus amplifiers (16-bit analog-to-digital conversion; 0.5-µV resolution; ±16.384-mV operating range; 5,000-Hz sampling; 0.1- to 250-Hz band pass) (Brain Products, Munich, Germany) and transmitted via fiber-optic cables. Data recording during the baseline night was made with BrainVision Recorder (v. 1.20) software while experimental nights data was readout in real-time, filtered 0.5-58 Hz, and relayed to custom MATLAB scripts for online EEG power analysis using OpenVibe software ^65^. EEG was digitized at an online sampling rate of 500 Hz and was referenced to the vertex. Electrode impedances were kept below 20 kΩ.

#### EEG Data Processing

Offline EEG data analysis was conducted with MATLAB (Mathworks Inc., Natick, MA) using the EEGLAB toolbox ^60^ and custom scripts. Sleep recordings were re-staged offline according to standard criteria to verify online sleep staging ^59^. EEG data was epoched at 60 seconds prior to each awakening, consistent with the time interval used for prediction (see *Procedure*). Twenty epochs were removed from analysis due to the presence of EEG artifacts or because sleep stage could not be confirmed. Data were bandpass filtered between 0.5 and 58 Hz using a Hamming windowed sinc FIR filter. Bad channels were identified by visual inspection and interpolated using spherical splines. Spectral power density was computed using Welch’s modified periodogram method. Source localization was performed in Brainstorm ^66^ with a noise-normalized estimate using standardized low-resolution brain electromagnetic tomography (sLORETA ^62^). Group-level analysis on EEG sources was conducted using the GLM framework implemented in SPM12 (Wellcome Trust Department of Imaging Neuroscience, University College London). Whole-cortex analyses were conducted and correction for multiple comparisons was performed using topological FDR ^67^(cluster forming threshold of *P* = 0.0005; cluster size threshold of *P* < 0.001; *k* = 100 vertices).

#### Procedure

Participants were invited to spend three consecutive nights in the lab. During the first night (baseline) they were allowed to sleep without interruption. Individual thresholds for low-frequency (1-4 Hz) and high-frequency (18-25 Hz) band power were computed from baseline night recordings by binning all NREM sleep into 60-second epochs and computing the top and bottom 15% thresholds from the distribution for each band for each participant over the entire baseline night (Fig. 5A, middle panel). On the experimental (second and third) nights, participants were awoken from NREM (sleep stage 2 or 3) when their neural activity surpassed a bispectral threshold in low- and high-frequency band power from an electrode cluster centered over the posterior hot zone. To ensure a stable sleep stage had been reached, participants were required to be in NREM for a minimum of 120 seconds before the algorithm initiated an awakening. The bispectral threshold required that for an DE prediction to be initiated low-frequency power had to be less than or equal to the lower 15% threshold while high-frequency power had to simultaneously meet the top 15% threshold for that subject. Likewise, for a NE prediction to be initiated low-frequency power had to be greater than or equal to the top 15% threshold while high-frequency power had to simultaneously be less than or equal to the bottom 15% threshold for that subject. This procedure was motivated by the results of experiments 1 and 2, and aimed to predict the presence of DE during time epochs in which the cortex was not only not bistable but also displayed correlates of neural firing (and vice versa for predictions of NE)^14,17^. Upon awakening participants completed a short version of the sleep questionnaire used in experiments 1 and 2.

### Selection of frequency bands and timeframes

#### Experiments 1 and 2

For low frequency power comparisons, we selected the 1-4 Hz band, as this frequency range is associated with the presence of slow waves, which have been linked to the loss of consciousness. Slow waves have been shown to exist not only in NREM sleep, but also in in REM sleep ^68^.

For high-frequency power comparisons, a slightly lower frequency range (20-50 Hz) was chosen for NREM sleep compared to REM sleep (25-50 Hz), given that NREM sleep is typically characterized by slower frequencies ^69^. Frequencies above 50 Hz were not included in order to avoid muscle and powerline artifacts. Frequencies below 20 Hz were avoided because of the potential overlap with fast sleep spindles, which can reach 18 Hz ^70^ ^71^.

#### Experiment 3

In experiment 3, the real time predictions were made based on an electrode cluster over posterior parietal-occipital cortex and this region is less prone to electrodermal artifacts. For this reason we designed this paradigm with a slightly less conservative low-frequency bandpower range (0.5-4.5 Hz) compared to experiments 1 and 2 (1-4 Hz), in order to capture and make use of the low end of low-frequency oscillatory activity toward predicting the presence or absence of experience.

In experiment 3, we selected the 18-25 Hz as the frequency range of interest in guiding real time predictions in NREM sleep to avoid confounds associated with eye movements, which have been shown to produce artifacts primarily in frequencies >25 Hz and particularly in the gamma band. In experiments 1 and 2 we had the advantage of being able to apply Independent Component Analysis (ICA) during offline post-processing of the data, which effectively suppresses such artifacts. This procedure could not be applied online in experiment 3.

#### Selection of timeframes

Since experience-related brain activity for the content of experiences was assumed to be very short-lived and subjects were specifically instructed to refer to the ‘very last’ experience before the alarm sound when reporting the presence of specific contents, we considered short timeframes before the awakening when performing content-specific contrasts, ranging from 0 to −8s with respect to the awakening. While this was relatively straightforward for the content, when the subjects had to tell whether they had had an experience or not, or could just not remember the content, it was more difficult for them to tie the answers to a specific timeframe. Therefore, we opted to include slightly longer timeframes (20s before the awakening) to account for possible variability when comparing DE, DEWR and NCE.

### Data availability

Data that are relevant to the publication are available upon request.

## Supplementary Material

**Supplementary Figure 1.**
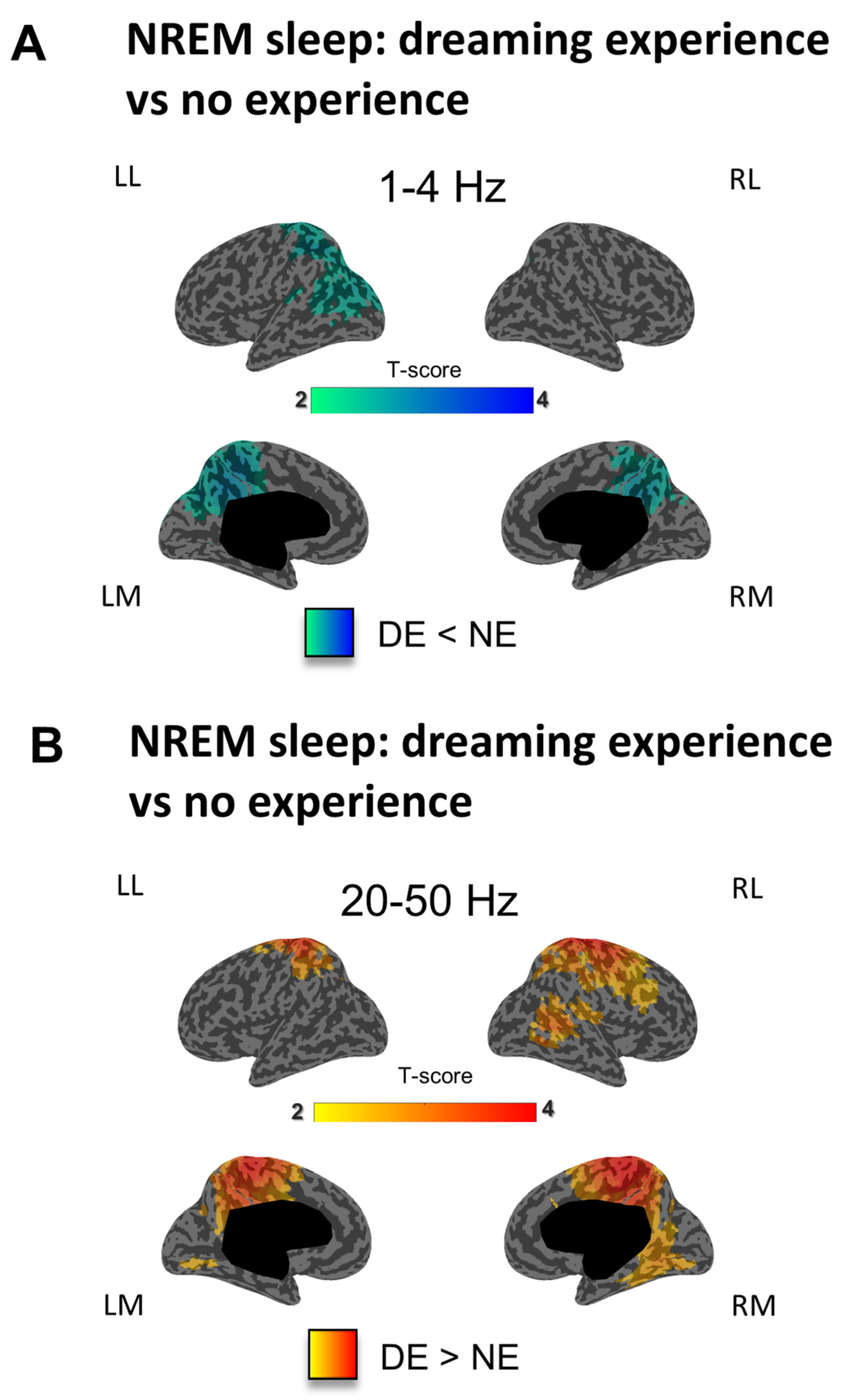
Dreaming experience vs. no experience (experiment 2). **A.** Inflated cortical maps illustrating the topographical distribution of t-values for the contrast between DEs and NEs at the source level for low-frequency power (1-4 Hz) in NREM sleep (last 20 seconds before the awakening) for experiment 2. Only significant differences at the p<0.05 level, obtained after correction for multiple comparisons are shown (two-tailed, paired t-tests, 7 subjects, t(6) ≥ 2.45). ***B***. Same as A for high-frequency power (25–50 Hz) in NREM sleep (two-tailed, paired t-tests, 7 subjects, t(6) ≥ 2.45).

**Supplementary Figure 2.**
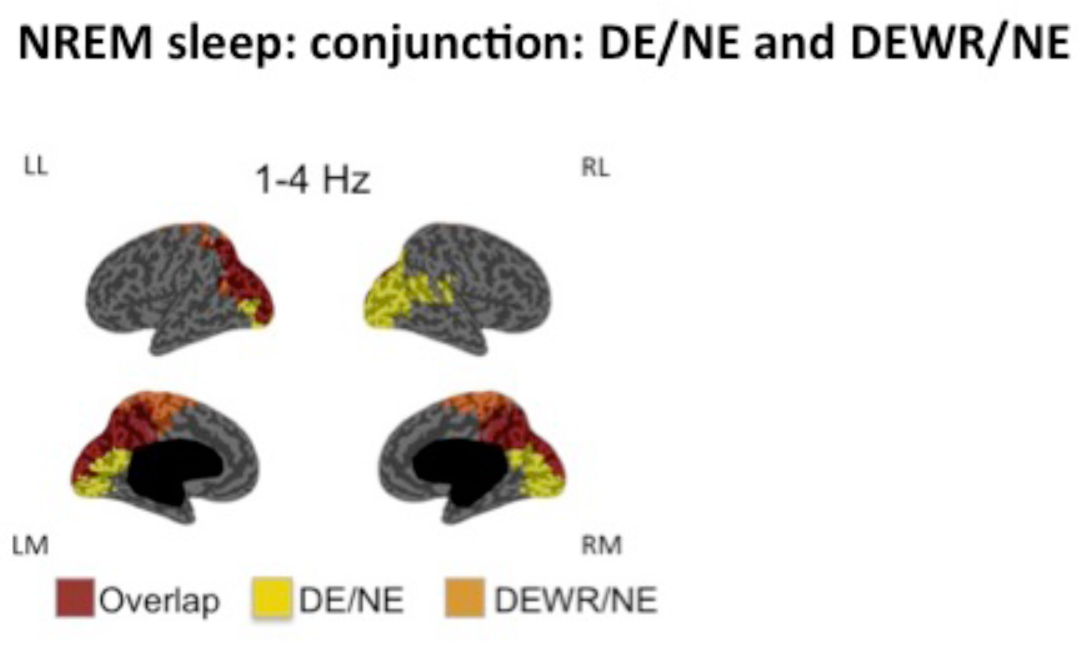
Conjunction between DE/NE and DEWR/NE contrasts. Conjunction maps showing the differences and overlap between the two contrasts (DE/NE and DEWR/NE) for low-frequency power in NREM sleep.

**Supplementary Figure 3.**
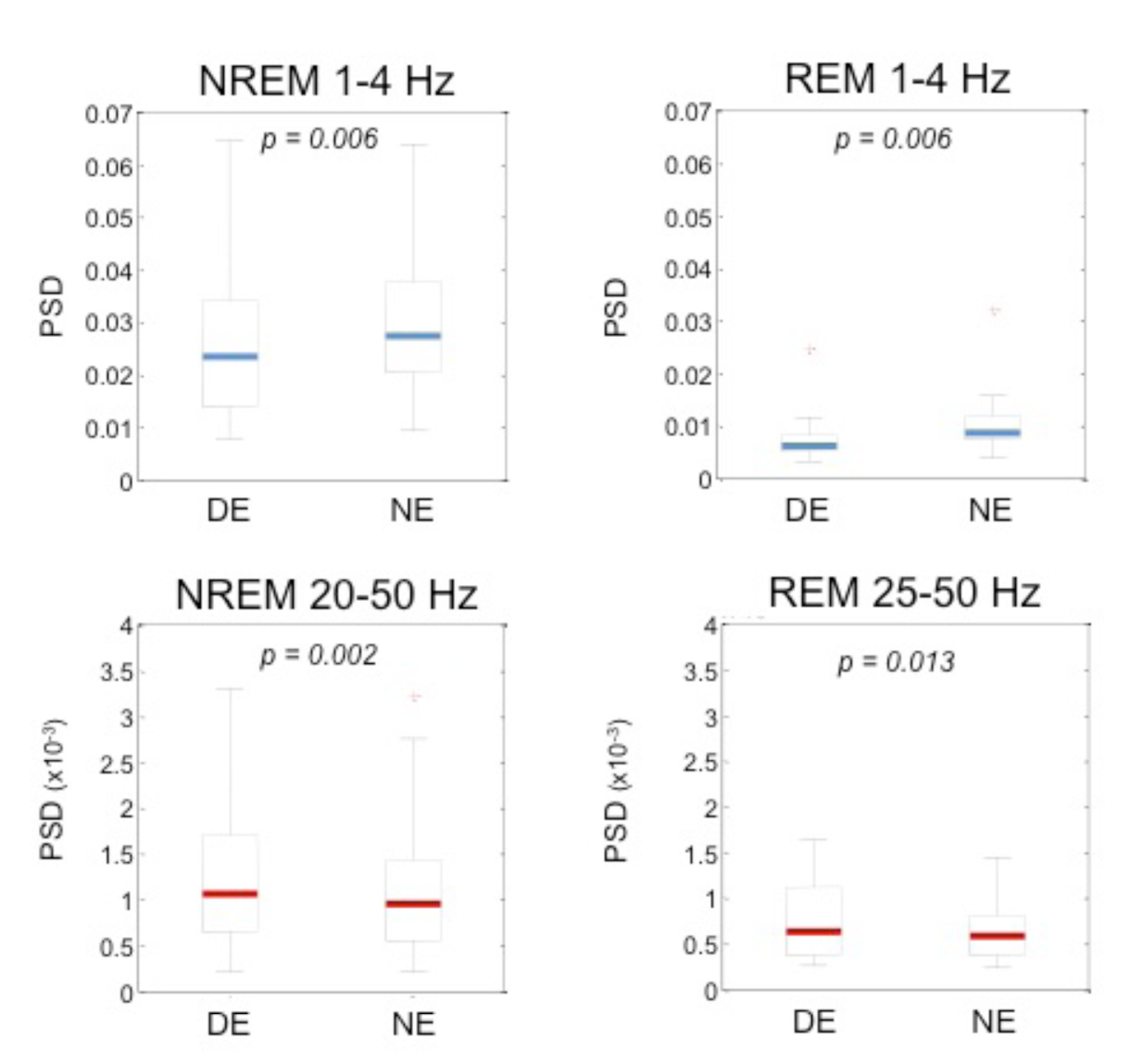
Absolute power values for DE and NE. Boxplots showing power spectral density (PSD) values for DE and NE, averaged over the posterior hot zone, (determined by the overlap between the DE/NE contrast in REM and NREM sleep, as shown in Figure 2B), for high- and low-frequency bands in NREM (n=32) and REM sleep (n=10). Significance was determined using two-tailed paired t-tests (NREM LF: t(31)= −2.98; NREM HF: t(31)= 3.46; REM LF: t(9)= −3.59; REM HF: t(9)= 3.10). Whiskers correspond to 3 standard deviations from the mean.

**Supplementary Table 1.** Examples of reports of most recent dreaming experiences.

| Stage | Time of night | Report |
| --- | --- | --- |
| N2 | 1:44 am | I was daydreaming. I was holding a musket, like a rifle, and I was skipping through images with it. Last image was a piece of pie, I am not sure but I think it was. |
| N2 | 5:23 am | I was thinking about perfume and fragrance. The very last word was 'fragrance'. |
| N2 | 5:55 am | The image of a Buddha belly, a bare belly. |
| N2 | 6:51 am | I saw a person waiting in a car. Maybe it was me. |
| N3 | 1:26 am | I was seeing geometric shapes that were moving very fast. |
| N3 | 5:18 am | I was trying to tell the difference between a homemade and a store-bought basket or tray of pastry. |
| N3 | 6:09 am | The last thing was raspberries, a pint of raspberries. |
| REM | 3:28 am | I was doing this experiment with another girl. I asked her what time it was and she said 7:07. No, she actually said 6:55. Her boyfriend was in the room, too. The last scene was just her face. It was quite a long dream before that. |
| REM | 3:59 am | I saw my brother eating hair on a plate. |
| REM | 6:05 am | It was the end of a movie. I was getting out on the street through a door. At that moment I hear the noise of the alarm sound and woke up. |
| REM | 6:36 am | In the last scene I was riding a bicycle in a street in B. Before that I was talking to someone in a court. Somebody taking care of the court. He gave lots of explanations about flowers. |

**Supplementary table 2.** Proportion of dream experiences in NREM and REM sleep.

| **Experiment 1 (NREM analysis, n=32, 1 night per subject)** |  |  |  |  |
| --- | --- | --- | --- | --- |
|  | DE | DEWR | NE | Total |
| NREM (N2) | 97 (42%) | 44 (19%) | 92 (39%) | 233 (100%) |
| **Experiment 1 (REM analysis, n=6, 1 night per subject) \*** |  |  |  |  |
|  | DE | DEWR | NE | Total |
| REM | 12 (55%) | 0 (0%) | 10 (45%) | 22 (100%) |
| **Experiment 2 (n=7, 5-10 nights per subject)** |  |  |  |  |
|  | DE | DEWR | NE | Total |
| NREM (N2 & N3) | 193 (33%) | 225 (38%) | 168 (29%) | 586 (100%) |
| REM | 131 (77%) | 31 (18%) | 8 (5%) | 170 (100%) |

\* The proportion of DE and NE in Experiment 1 reflects the criterion of only including subjects that have both DE and NE during the night.

**Supplementary table 3.** Prediction accuracy of DE and NE in sleep.

| Participant | DE ACC | NE ACC | N awakenings | 0.5-4.5 Hz thresh | 18-25 Hz thresh |
| --- | --- | --- | --- | --- | --- |
| 1 | 0.89 | 0.75 | 14 | [30.0 217.0] | [0.25 0.40] |
| 2 | 0.89 | 0.75 | 13 | [30.0 165.0] | [0.11 0.22] |
| 3 | 1.0 | 1.0 | 8 | [32.0 188.0] | [0.09 0.30] |
| 4 | 1.0 | 0.71 | 16 | [42.9 153.2] | [0.19 0.43] |
| 5 | 0.89 | 0.86 | 16 | [15.0 94.0] | [0.08 0.20] |
| 6 | 0.78 | 1.0 | 10 | [14.8 63.0] | [0.06 0.11] |
| 7 | 1.0 | 1.0 | 7 | [19.0 154.0] | [0.09 0.28] |

**Supplementary table 4.** High/Low frequency power ratio DE > NE.

| Region | Volume<br>(k<sub>E</sub>) | Peak<br>*t*-value | Peak MNI |  |  |
| --- | --- | --- | --- | --- | --- |
|  |  |  | X | Y | Z |
| R precuneus/cuneus/SPL/SOC | 2920 | 11.84 | 19 | -63 | 58 |
| L precuneus/cuneus/STS/SOC | 1056 | 11.09 | -7 | -79 | 51 |
| R MFG | 309 | 10.78 | 37 | 50 | -6 |
| R lingual gyrus | 158 | 9.39 | 20 | -57 | -13 |
| R precentral gyrus | 153 | 9.08 | 42 | -8 | 65 |
| L IFG | 162 | 7.98 | -53 | 24 | -6 |

\* All clusters significant at $p < 0.001$ , FDR cluster corrected.
R: right; L: left; SPL: superior parietal lobule; STS: superior temporal sulcus;
SOC: superior occipital cortex; MFG: Middle frontal gyrus; IFG: inferior frontal gyrus

**Supplementary table 5.**
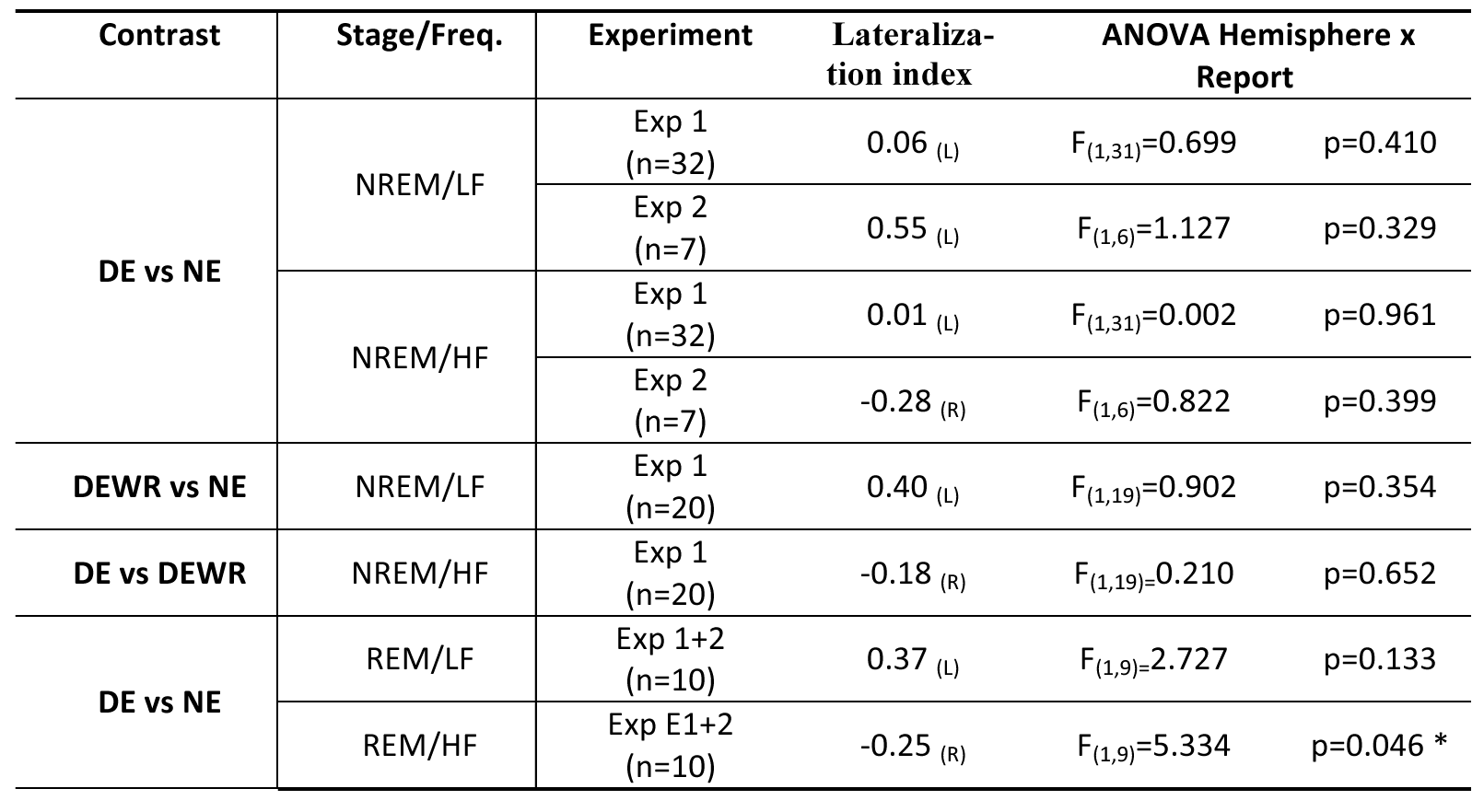
lateralization indices. Lateralization of findings. For each contrast, the brain hemisphere including the relative majority of activated voxels was identified (Lateralization Index, LI)^64^ and repeated measure ANOVA was performed to evaluate the possible interaction between hemisphere and type of report. * p<0.05.

## End notes

**Supplementary Material** is linked to the online version of the paper at www.nature.com/nature.

## Acknowledgments

Supported by NIH/NCCAM P01AT004952 (GT), NIH/NIMH 5P20MH077967 (GT), Swiss National Science Foundation Grants 139778 (FS), 145571(FS) and 155120 (LP), Swiss Foundation for Medical Biological Grants 151743 and 145763 (FS), NIH/NINDS F32NS089348 (BB), UW Medical Scientist Training Program Grant T32 GM008692 (JJL), NIH Grant MH064498 (BRP).

The authors thank David Bachhuber, Emmanuel Carrera, Anna Castelnovo, Amelia Cayo, Chiara Cirelli, Chadd Funk, Matthew Gevelinger, Jeanne Harris, Armand Mensen, Poorang Nori, Richard Smith, Laurène Vuillaume, Sophy Yu, Corinna Zennig and the undergraduate research assistants for help with data collection, sleep scoring, technical assistance and helpful discussions.

## Author Contributions

FS, LP, BB, JL, MB, BR, BP and GT designed the experiments, FS, LP, BB and JL conducted the experiments, FS, LP, BB and GB analyzed the data, FS, LP, BB and GT wrote the paper.

## Author Information

GT and BR are involved in a research study in humans supported by Philips Respironics. This study is not related to the work presented in the current manuscript. The other authors have indicated no financial conflicts of interest. Correspondence and requests for materials should be addressed to GT.

